# Structural and biophysical characterization of HNF-1A as a tool to study MODY3 diabetes variants

**DOI:** 10.1101/2021.12.20.473529

**Authors:** Laura Kind, Arne Raasakka, Janne Molnes, Ingvild Aukrust, Lise Bjørkhaug, Pål Rasmus Njølstad, Petri Kursula, Thomas Arnesen

## Abstract

Hepatocyte nuclear factor 1A (HNF-1A) is a transcription factor expressed in several embryonic and adult tissues, modulating expression of numerous target genes. Pathogenic variants in the *HNF1A* gene cause maturity-onset diabetes of the young 3 (MODY3 or *HNF1A* MODY), characterized by dominant inheritance, age of onset before 25-35 years of age, and pancreatic β-cell dysfunction. A precise diagnosis alters management as insulin can be exchanged with sulfonylurea tablets and genetic counselling differs from polygenic forms of diabetes. More knowledge on mechanisms of HNF-1A function and the level of pathogenicity of the numerous *HNF1A* variants identified by exome sequencing is required for precise diagnostics. Here, we have structurally and biophysically characterized an HNF-1A protein containing both the DNA binding domain and the dimerization domain. We also present a novel approach to characterize HNF-1A variants. The folding and DNA binding capacity of two established MODY3 HNF-1A variant proteins (P112L, R263C) and one variant of unknown significance (N266S) were determined. All three variants showed reduced functionality compared to the wild-type protein. While the R263C and N266S variants displayed reduced binding to an HNF-1A target promoter, the P112L variant was unstable *in vitro* and in cells. Our results support and mechanistically explain disease causality for all investigated variants and allow for the dissection of structurally unstable and DNA binding defective variants. This points towards structural and biochemical investigation of HNF-1A being a valuable aid in reliable variant classification needed for precision diagnostics and management.

## Introduction

Maturity-onset diabetes of the young (MODY) is a monogenic, autosomal-dominant form of diabetes mellitus, which is characterized by impaired β-cell function and insulin regulation. The disease is caused by mutations in one of 11 genes crucial for pancreatic β-cell development and homeostasis (1). MODY3, which contributes to ~40-70% of all MODY cases, is associated with pathogenic variants of the hepatocyte nuclear factor 1A (*HNF1A*) gene (2). The encoded transcription factor HNF-1A is expressed both in the embryonic and adult stages of the liver, gall bladder, pancreas, gastro-intestinal tract, kidney, urinary bladder, as well as the adult bone marrow and immune system (3). This wide expression pattern of HNF-1A indicates diverse molecular functions in the human body. Specifically in the endocrine pancreas, HNF-1A regulates the expression of numerous target genes involved in glucose regulation and metabolism, such as those encoding the insulin receptor, HNF-4A, the glucose transporter GLUT2, and glucose-6-phosphatase (4–6). In a genome location experiment using human pancreatic islets, HNF-1A was found to occupy promoter regions of 106 islet genes (4) and HNF-1A deficiency in mice resulted in broad changes of islet gene expression patterns (5). Studies on the molecular structure and function of this important transcription factor is likely to improve our understanding of pancreatic islet biology and underlying mechanisms for the development of MODY3.

HNF-1A is a multi-domain protein (Fig. 1A), which contains an N-terminal dimerization domain (DD), a central DNA binding domain (DBD), and a C-terminal transactivation domain (TAD) (3). Three isoforms (A, B, C) have been described, which differ in the length of the C-terminal TAD and are differentially expressed in fetal and adult tissues (7). The N-terminal DD is a 32-residue α-helical region, which can fold and dimerize independently from the residual domains (8,9). The DD forms a heterotetramer with the pterin-4 alpha-carbinolamine dehydratase 1 (PCBD1), which stabilizes the dimeric form of HNF-1A (10,11). A 50-residue long linker separates the DD from the DBD, which consists of a Pit-Oct-Unc-specific (POU_S_) domain and a homeodomain (POU_H_) and can bind to the inverted palindromic consensus DNA sequence GTTAATNATTAAC (12). The 200-residue long DBD contains a nuclear-localization signal (NLS) preceding the POU_H_ domain, ensuring translocation of HNF-1A into the nucleus (13). The structure of the isolated POU_H_ domain from rat has been solved by X-ray crystallography and nuclear magnetic resonance spectroscopy, revealing an atypical homeodomain with an extended loop between the α2- and α3-helices (14,15). Chi *et al*. reported the crystal structure of the entire DBD bound to a high-affinity promoter, corresponding to the consensus sequence for optimal HNF-1A binding (16). The POU_S_ and POU_H_ domains form a stable interface when bound to the DNA, potentially increasing affinity and stability of the complex (16). In addition, HNF-1A contains a 450-residue long TAD in the C-terminus, which is involved in gene transcription activation and likely interacts with numerous transcription factors, co-activators, and repressors (13,17,18). Apart from structural models for the isolated DD and DBD domains, the HNF-1A protein remains structurally and biophysically uncharacterized.

**Fig. 1.**
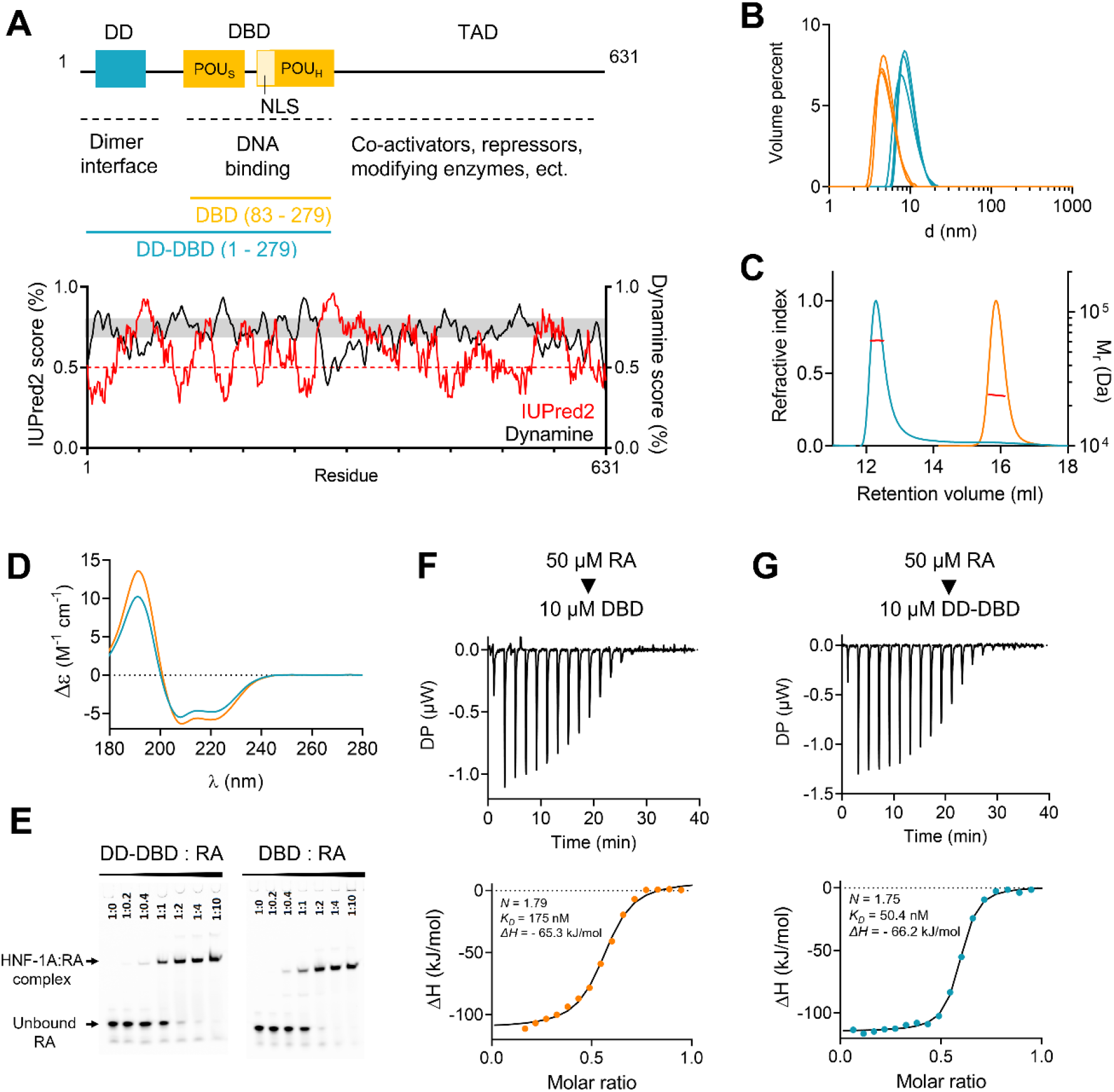
Biophysical and functional characterization of HNF-1A DBD and DD-DBD constructs. A) Top: Domain overview of HNF-1A with residue numbers and indicated domain functions. DD-DBD (residues 1 – 279) and DBD (residues 83-279) constructs used for *in vitro* protein characterization are shown in turquois and orange, respectively. Bottom: IUPred2 and DynaMine prediction scores for full-length HNF-1A. IUPred2 scores above 0.5 indicate propensity for disorder, while scores below 0.5 predict ordered regions (26). DynaMine scores below 0.7 indicate disorder, while scores above 0.8 predict order. DynaMine prediction in the grey zone (scores between 0.7 and 0.8) indicate residues with highly context-dependent dynamics (27). B) DLS of DBD and DD-DBD. C) SEC-MALS profiles for DBD and DD-DBD. Red line: molecular weight based on MALS and refractive index. D) SRCD spectra for DBD and DD-DBD. E) EMSA titration native gels showing complex formation of RA and respective HNF-1A construct. F, G) ITC titration curves for DBD and DD-DBD, respectively. Titrant: 50 μM double-stranded RA oligonucleotide. Titrand: 10 μM DBD (F) or DD-DBD (G).

According to the ClinVar database (19), 418 variants of the *HNF1A* gene have been reported, out of which 131 are recognized as pathogenic or likely pathogenic. Remaining variants are classified as likely benign, benign, or variants of uncertain significance (VUS). Diabetes-associated missense mutations occur at highest rates in the DBD, while the mutation rate is intermediate in the DD and low in the TAD (16). In this study, we aimed to structurally and functionally characterize the entire DD-DBD HNF-1A region, harboring most of the variants causing MODY3, and to establish the system as a tool to evaluate the dysfunctional characteristics of MODY3-associated variants *in vitro*. To this end, we compared recombinantly-expressed and purified wild-type (WT) protein and two established pathogenic MODY3 variants: p.Pro112Leu and p.Arg263Cys, hereafter referred to as P112L and R263C (20,21). P112L and R263C, respectively, contribute to approximately 5.7% and 1.4% of all MODY3 probands registered in the Norwegian MODY registry to date. Both variants have previously been characterized regarding cellular functions, but molecular mechanisms remain enigmatic (20,22,23). Our biophysical *in vitro* approach revealed defects regarding protein folding and DNA binding ability, providing explanations for the pathogenicity of these established MODY3 variants. Finally, we used this model system to investigate the *HNF1A* p.Asn266Ser (N266S) variant, which is currently classified as a VUS. N266S is considered as a rare variant, with only one registered proband/family within the Norwegian MODY registry to date. This variant was not described functionally previously, and our biophysical characterizations revealed defects at the protein level, supporting disease causality for the N266S variant.

## Results

### Biophysical and functional characterization of two HNF-1A constructs

The individual DD and DBD of HNF-1A can be recombinantly expressed and purified for structural studies (8–10,16). In contrast, the C-terminal TAD has not been studied using recombinant expression systems, which is likely due to the disordered nature typical for TADs (24,25). Indeed, using disorder predictions we confirmed that large regions in the TAD have a high tendency for disorder (Fig. 1A). To ensure soluble protein expression, we therefore omitted the TAD in this study and created two HNF-1A constructs: HNF-1A 83-279 contained the isolated DBD, while HNF-1A 1-279 contained the DD, a linker, and the DBD. These constructs are hereafter referred to as HNF-1A DBD and HNF-1A DD-DBD, respectively (Fig. 1A).

Both protein constructs were purified to homogeneity. DBD and DD-DBD exhibited a slower SDS-PAGE migration than expected based on theoretical molecular weight (Fig. S1); however, the protein identity could be confirmed by peptide mass fingerprinting. Initial dynamic light scattering (DLS) experiments indicated a hydrodynamic radius of approximately 6.0 nm and 10.7 nm for construct DBD and DD-DBD, respectively (Fig. 1B). These observations were according to expectations, as construct DD-DBD had a higher theoretical mass than DBD and was expected to be dimeric in solution. Size-exclusion chromatography coupled to multi-angle light scattering (SEC-MALS) confirmed this result (Fig. 1C), as construct DBD exhibited a molecular weight of 24.0 ± 2.4 kDa (monomeric theoretical weight: 22.8 kDa), and construct DD-DBD showed a molecular weight of 61.3 ± 3.7 kDa (monomeric theoretical weight: 31.2 kDa, dimeric theoretical weight: 62.4 kDa).

Synchrotron radiation circular dichroism (SRCD) measurements were conducted to investigate the secondary structure of DBD and DD-DBD, which adopted an α-helical fold. DBD gave a stronger α-helical signal, likely due to the absence of the disordered linker present in the DD-DBD construct (Fig. 1D). These findings were in line with previous reports regarding HNF-1A structure and were important quality-control steps, ensuring that both constructs were suitable for subsequent studies of MODY3 variants.

We qualitatively assessed DNA binding ability of DBD and DD-DBD by electrophoretic mobility shift assays (EMSA), in which the purified proteins were added in different ratios to a double-stranded rat albumin promoter oligonucleotide (RA), and the migration behavior of RA was observed during native gel electrophoresis. As the free oligonucleotide probe migrates faster than when in a macromolecular complex, EMSA allows the separation of bound and unbound oligonucleotide and to approximate binding affinities between a protein and the specific promoter oligonucleotide. EMSA showed that both DBD and DD-DBD bound RA and that the protein:RA complexes differed in size, accordingly to the described size differences of the purified HNF-1A constructs (Fig. 1E). Complex formation started to appear at a molar RA:protein ratio of 1:0.4 and seemed to be nearly complete at a molar ratio of 1:2.

We quantified DNA binding properties by isothermal titration calorimetry (ITC), from which we extracted thermodynamic parameters of complex formation (Fig. 1F,G, Table 2). The stoichiometry values from triplicate measurements were determined to 1.97 for construct DD-DBD and 1.89 for construct DBD (Table 2), which corresponds to a RA:protein binding stoichiometry of 1:2. This observation is in line with the crystal structure of an RA:DBD complex (PDB ID: 1IC8; (16)), in which two DBD molecules bind to one double-stranded RA oligonucleotide. Enthalpy changes (*ΔH*) for both constructs were in the same range (Table 2), suggesting a similar binding mechanism. The affinity between the protein and RA was high, as the average equilibrium dissociation constant *K_D_* was determined to 100 nM for DD-DBD and 114 nM for DBD (Table 2). EMSA and ITC measurements demonstrated that both constructs were able to bind DNA, establishing them as suitable models for *in vitro* characterization of disease-causing variants impacting DNA binding.

### HNF-1A DBD and DD-DBD exhibit an extended conformation in solution

Since both DBD and DD-DBD were predicted to contain flexible linker regions (Fig. 1A), we performed small-angle X-ray scattering (SAXS) to obtain information about the conformation and flexibility of the proteins (Fig. 2). The scattering curves of DBD and DD-DBD gave a first indication that the proteins differ in molecular shape (Fig. 2A). The maximal molecular dimension (*D_max_*) could be extracted from the distance distribution function (Fig. 2B): 10.2 nm for DBD and 24.4 nm for DD-DBD. These results agree with our DLS and SEC-MALS data (Fig. 1B,C). Kratky analysis of the SAXS data showed that both constructs adopt an extended, non-globular conformation, which can be explained by the linkers between the POU_S_ and POU_H_ domain, as well as an unfolded region between the DD and DBD (Fig. 2C). Indeed, a CRYSOL analysis, comparing the DNA-bound state of DBD (PDB ID: 1IC8) and the free state in solution, revealed a difference in compactness and overall shape (Fig. 2A). An *ab initio* model was generated for DBD and superposed with the DBD crystal structure from the DBD:RA complex (16), which illustrates the difference in molecular conformation (Fig. 2D). It appears that the stable interface between the POU_S_ and POU_H_ domains reported by Chi *et al*. only forms upon DNA binding. A rigid body model with built linker regions also suggested that the POU_S_ and POU_H_ domains do not form intimate contacts in the unbound state (Fig. 2E). In order to probe for changes in secondary structure content upon DNA binding, we performed SRCD experiments for both DBD alone and a DBD:RA complex (Fig. 2F). We concluded from the highly similar spectra that there is no notable change in secondary structure content during complex formation, but that the 18-residue linker between the POU_S_ and POU_H_ domains becomes more compact upon DNA binding. An *ab initio* model (Fig. 2G) and a rigid body model (Fig. 2H) were generated for construct DD-DBD and revealed a similar behavior. The protein adopts an elongated, extended shape in solution.

**Fig. 2.**
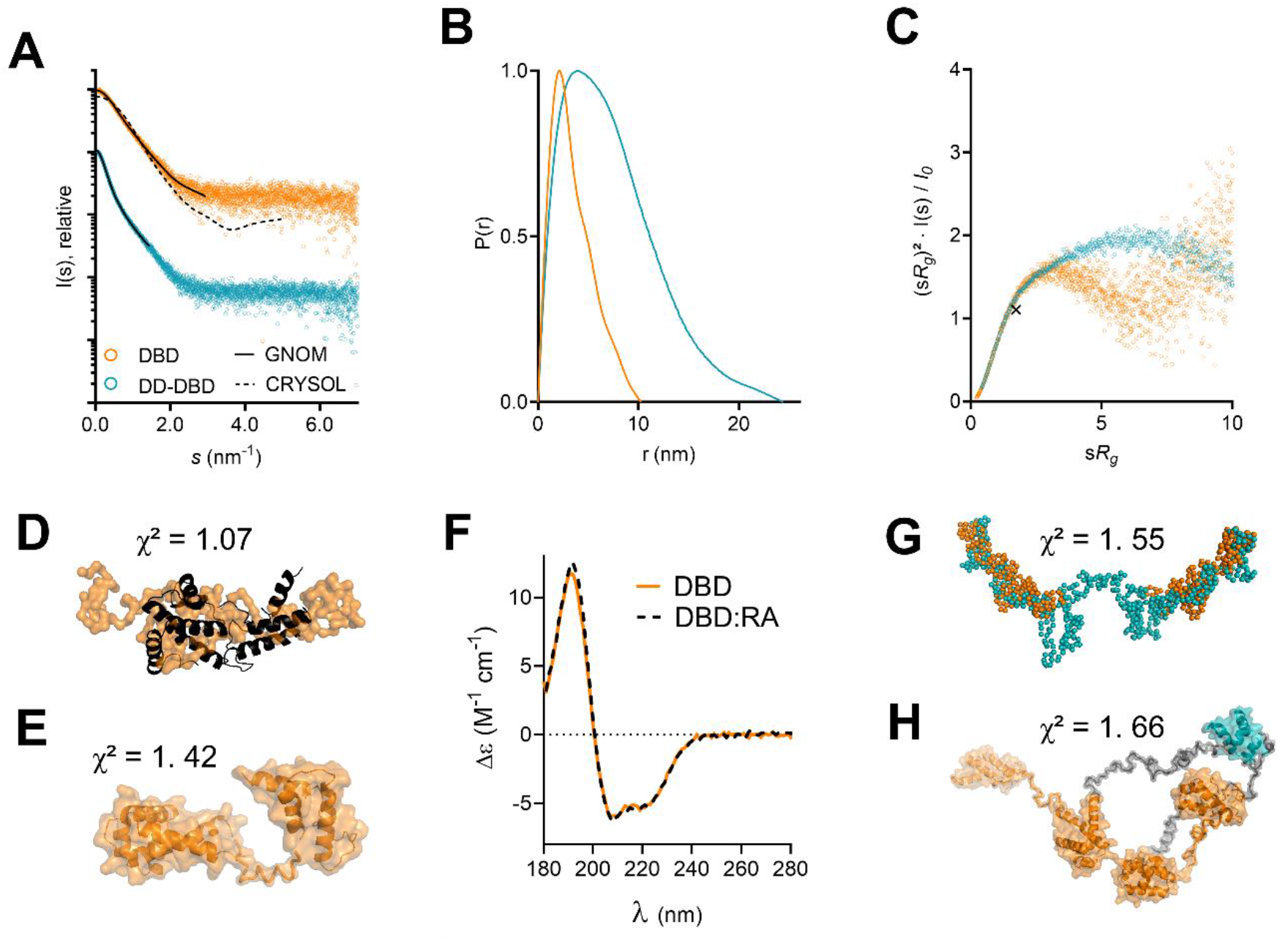
SAXS data for DBD and DD-DBD. A) Scattering curves for DBD and DD-DBD. Solid line: GNOM fit. Dashed line: CRYSOL fit with crystal structure model from PDB 1IC8 (16) (χ^2^ = 10.4). B) Distance distribution functions for DBD and DD-DBD. C) Dimensionless Kratky plot for DBD and DD-DBD, with the cross indicating the expected maximum for a globular particle (√3, 1.104) (28). D) GASBOR model of DBD (orange), superposed with the oligo-bound model from DBD:RA crystal structure (16). E) CORAL model for DBD. F) SRCD spectra for DBD in presence and absence of RA. G) GASBOR model for DD-DBD (teal), overlaid with GASBOR model of DBD (orange) at estimated positions. H) CORAL model of DD-DBD. DBD residues are shown in orange, linker residues in grey, and DD residues in blue.

### HNF-1A P112L shows severely reduced solubility *in vitro* and decreased stability in cells

Following the biophysical and biochemical characterization of the two HNF-1A WT constructs, we aimed to utilize this system to study diabetes-causing variants of HNF-1A. We chose to explore two established pathogenic variants (P112L and R263C) and one VUS (N266S) to elucidate disease mechanisms of MODY3. All three mutation sites are located in the DBD of HNF-1A (Fig. 3A–D). The alignment of HNF-1A sequences from model organisms (residues 1 – 279) showed that the three affected residues are highly conserved across species down to zebrafish and suggested low tolerance for sequence variance in regard to protein function (Fig. 3A). Pro112 is in the POU_S_ domain at the beginning of helix α2 (Fig. 3B,C). Considering the helix-terminating nature of proline residues, this position suggests that Pro112 might have an important role in the secondary structure formation of POU_S_. Both Arg263 and Asn266 are located in the POU_H_ domain (Fig. 3B). Both residues are found in the C-terminal helix α9, which is inserted into the major groove of the DNA (Fig. 3D). Arg263 partakes in ionic interactions with the backbone phosphate group of the adenine in the sixteenth position of the RA oligonucleotide (o.A16). In addition, Arg263 forms polar contacts with the backbone carbonyl group of Leu258 and possibly with the side chain of Thr260. Asn266 is situated in proximity of the DNA molecule and may be involved in DNA recognition. The side chain of Asn266 does not partake in direct contacts with other residues, but the polar nature and chain length of asparagine may be important for energetically favorable protein folding.

**Fig. 3.**
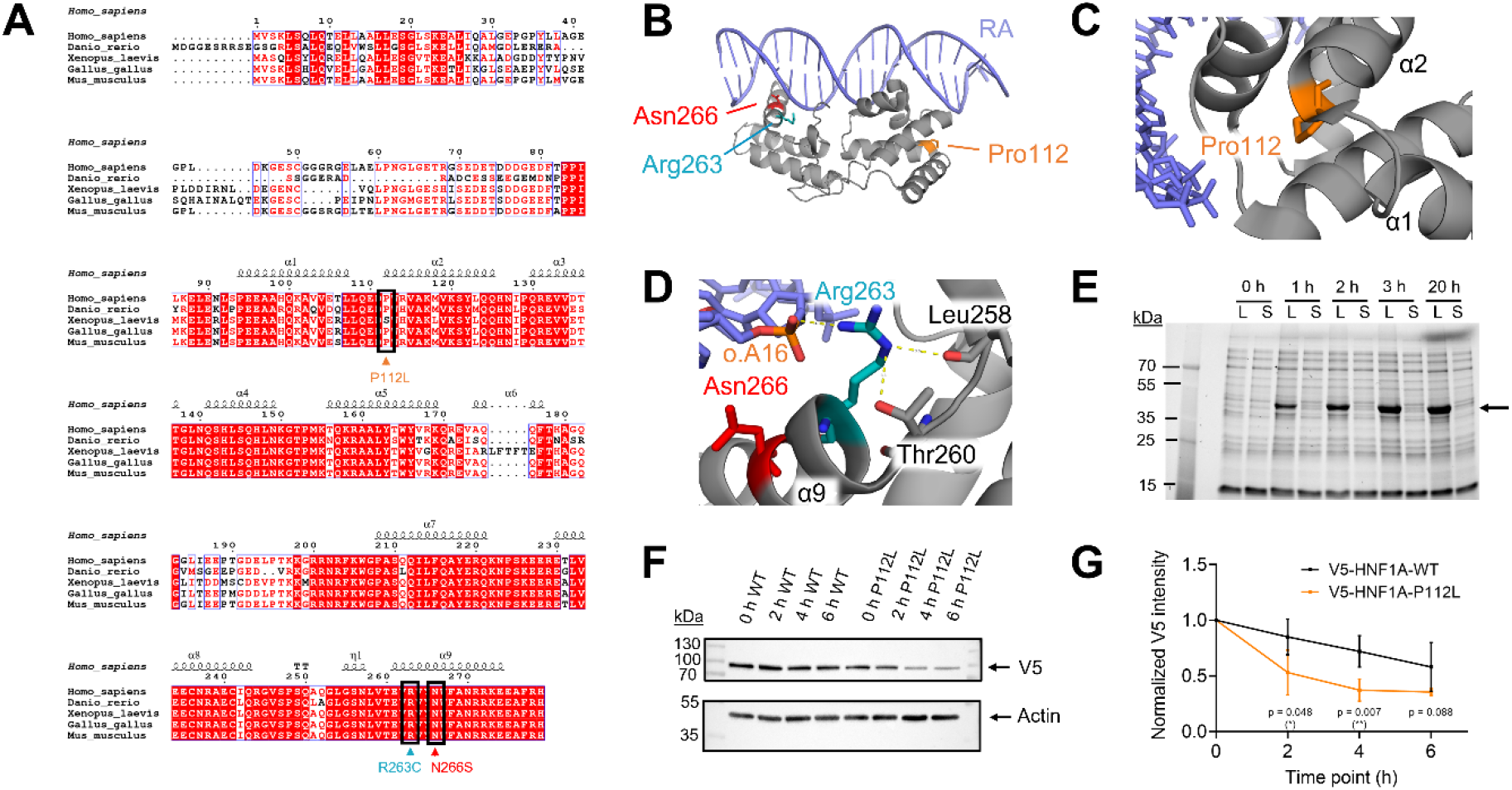
Overview of the studied variants and protein turnover dynamics of HNF-1A P112L. A) Multiple sequence alignment of HNF-1A sequences from model organisms, with variants of interest indicated by black boxes and arrows. B) Crystal structure of RA-bound DBD (PDB ID: 1IC8) with mutation sites highlighted. C) Magnified view on Pro112 and arrangement of neighboring residues. D) Magnified view on Arg263 and Asn266. o.A16 and neighboring residues Leu258 and Thr260 are indicated. E) Expression and solubility test for DD-DBD P112L (31.2 kDa) in *E. coli* Rosetta(DE3). Lysate (L) and soluble fraction (S) at different time points of expression (1 – 3 h: 37 °C, 20 h: 20 °C, 0 h: uninduced control). Arrow indicates over-expressed protein band with an apparent slow migration behavior, as observed for DD-DBD WT. F) Representative western blot membranes of CHX chase assay analysis of full-length WT and P112L protein turnover. G) Quantification of CHX chase assay for HNF-1A WT and P112L, based on normalization to whole protein amount from stain-free SDS-PAGE gel (N = 4, Fig. S2).

Initial expression tests in *E. coli* Rosetta(DE3) distinguished the P112L variant from the R263C and N266S variants. While the latter were equally soluble as the WT constructs, the P112L variant showed a severe reduction in protein solubility (Fig. 3E, Fig. S1A,B). Different lysis buffers were tested, varying pH, NaCl concentration, and amounts of stabilizing additives, but no solubilizing condition was found (data not shown). An additional screen in *E. coli* Lemo21(DE3) was conducted, in which protein expression levels can be tuned by titration of L-rhamnose into the expression media, leading to slower and more controlled protein expression. However, the P112L variant appeared insoluble, independent of the titrant amount (Fig. S1C). This observed insolubility of P112L indicated misfolding of the protein, leading to aggregation or incorporation into bacterial inclusion bodies.

To test the hypothesis of protein instability further, we performed protein turnover experiments in HeLa cells, employing a cycloheximide (CHX) chase assay (Fig. 3F,G, Fig. S2). The drug CHX inhibits protein synthesis in the cells, allowing the assessment of protein degradation dynamics. Using a specific anti-V5 antibody, the amount of full-length V5-WT and V5-P112L protein was detected at different time points of CHX treatment (Fig. 3F). Based on the V5-signal at time point 0 h, it was apparent that the P112L variant is expressed at lower levels than the WT protein at steady state conditions (Fig. 3F). Alternatively, the weaker V5-signals for P112L could arise from lower solubility of the P112L variant as compared to WT HNF-1A. The time-course of normalized protein levels indicated that the protein turnover of the P112L variant was elevated compared to the WT protein (Fig. 3G).

### HNF-1A R263C and N266S display normal protein folding and partially altered thermal stability

The established pathogenic variant R263C and the VUS N266S were expressed and purified like the WT constructs. We performed DLS experiments to investigate hydrodynamic diameters of the proteins and found that the variant constructs were similar to the respective WT constructs (Fig. 4A). SEC-MALS measurements confirmed that DBD/R263C was monomeric in solution (M_r_ = 23.9 ± 2.4 kDa), while DD-DBD/R263C formed a dimer in solution (M_r_ = 60.2 ±3.4 kDa) (Fig. 4B, Table S1). The corresponding N266S constructs were found to be in the same oligomeric state (DD-DBD/N266S: M_r_ = 61.9 ± 3.4 kDa, DBD/N266S: M_r_ = 24.1 ± 2.1 kDa). All variant elution profiles overlapped entirely with the respective WT SEC profile (Fig. 4B).

**Fig. 4.**
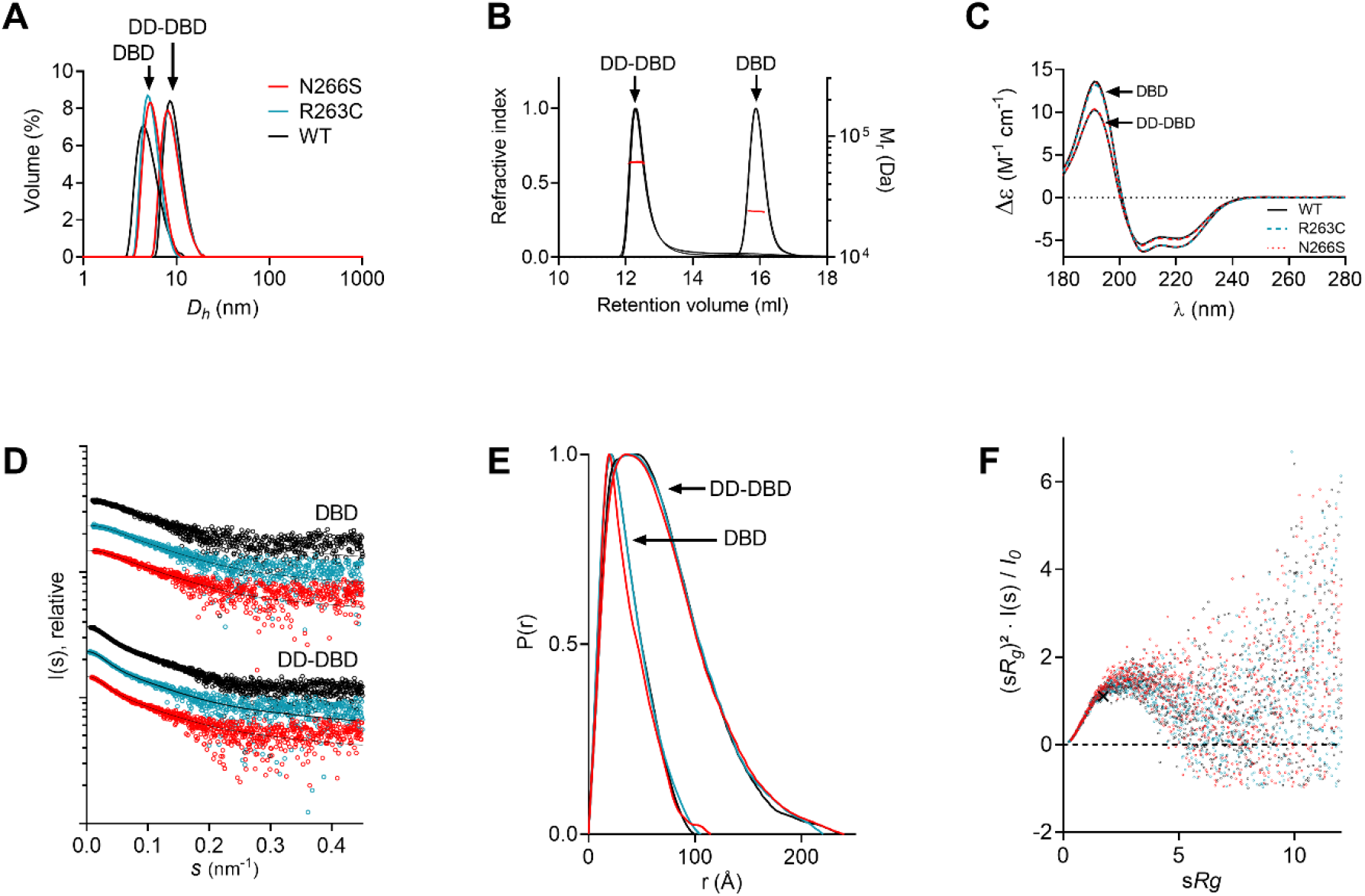
Biophysical characterization of HNF-1A R263C and N266S. A) DLS curves for DBD and DD-DBD variants. B) SEC-MALS elution profiles for DBD and DD-DBD variants. Red line: molecular weight based on MALS and refractive index. C) SRCD spectra for DBD and DD-DBD variants. D) SAXS scattering curves for DBD and DD-DBD variants. E) SAXS distance distribution functions for DBD and DD-DBD variants. F) Dimensionless Kratky plot for DBD variants, with the cross indicating the expected maximum for a globular particle (√3, 1.104) (28).

SRCD spectra of the DBD and DD-DBD variants overlaid with the corresponding WT spectra across the entire wavelength range, indicating identical secondary structure (Fig. 4C). We conducted SAXS experiments to investigate the solution structure of the studied HNF-1A variants (Fig. 4D–F, Table S3). The R263C mutation did not introduce changes in molecular dimension, molecular shape, or flexibility. In contrast, the N266S variant appeared slightly less compact, as a minor increase in *D*_max_ (DBD/WT: 10.0 nm, DBD/N266S: 11.5 nm) became apparent from the distance distribution function (Fig. 4E, Table S3).

Thermal stability was assessed by differential scanning fluorimetry (DSF), from which the melting temperature (*T_M_*) of each variant was extracted (Table 1, Fig. S4). For the WT constructs, the *T_M_* values were generally very low (DBD: 42.2 °C, DD-DBD: 35.8 °C), which likely results from the presence of the small, separated POU_H_ and POU_S_ domains, as well as the long, disordered linker in DD-DBD. The DSF protein stability assay also revealed that the (DD-)DBD/R263C variants were as stable as the respective WT constructs. In contrast, the N266S mutation was found to de-stabilize the protein during thermal denaturation, as the DBD/N266S variant exhibited a *T_M_* value that was 4 °C lower than the *T_M_* of the DBD WT construct (Table 1). The thermal instability of DBD/N266S may be connected to the less compact fold of the same construct, as observed in SAXS.

**Table 1.**
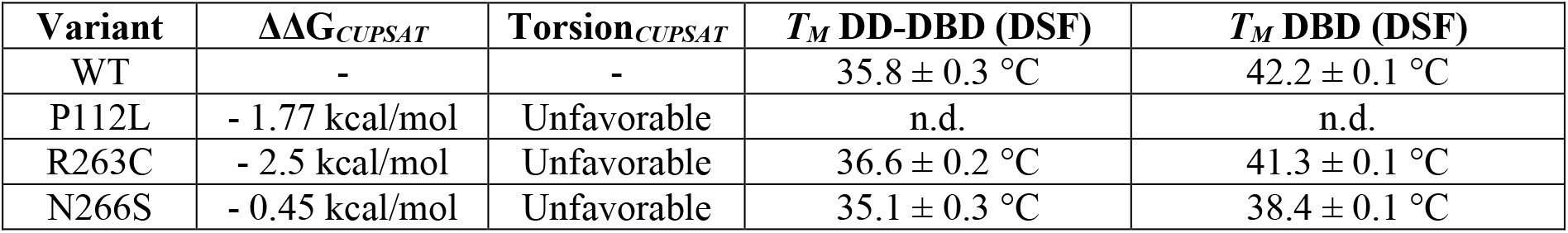
Thermal protein stability predictions by *CUPSAT* and experimental *T_M_* values determined by DSF analysis (mean ± SD; N = 4). n.d. – not determined.

| Variant | $\Delta\Delta G_{CUPSAT}$ | Torsion<sub>CUPSAT</sub> | $T_M$ DD-DBD (DSF) | $T_M$ DBD (DSF) |
| --- | --- | --- | --- | --- |
| WT | - | - | $35.8 \pm 0.3$ °C | $42.2 \pm 0.1$ °C |
| P112L | - 1.77 kcal/mol | Unfavorable | n.d. | n.d. |
| R263C | - 2.5 kcal/mol | Unfavorable | $36.6 \pm 0.2$ °C | $41.3 \pm 0.1$ °C |
| N266S | - 0.45 kcal/mol | Unfavorable | $35.1 \pm 0.3$ °C | $38.4 \pm 0.1$ °C |

We compared the experimentally determined stability differences with computational predictions by using the *CUPSAT* prediction algorithm (Table 1). *CUPSAT* was used to assess possible changes in free energy of protein folding (*ΔΔG*) upon point mutation, considering amino acid atom potentials and torsion angle distribution (29). All assessed variants (P112L, R263C, N266S) were predicted to introduce unfavorable torsion angles and to de-stabilize the protein structure (*ΔΔG* < 0), which likely contributes to the observed defects.

### HNF-1A R263C and N266S exhibit strongly reduced DNA binding ability

While the overall folding of R263C and N266S constructs appeared normal, the DNA binding ability was strongly affected (Fig. 5). We performed a dose-dependent EMSA to qualitatively measure DNA binding affinity (Fig. 5A,B). A faint complex band was observed at a 1:0.2 RA:protein molar ratio for WT DD-DBD, while a first complex band for DD-DBD/R263C was only visible at a molar RA:DD-DBD/R263C ratio of 1:0.4 (Fig. 5A). Full RA:DD-DBD binding was achieved at a ratio of 1:4, while the R263C variant did not reach complete binding even at a molar ratio of 1:10. The binding behavior for the R263C variant appeared shifted to the right side of the EMSA titration (Fig. 5A). The EMSA titration for DD-DBD/N266S also revealed a weaker protein DNA complex formation, although not as severe as for DD-DBD/R263C (Fig. 5A,B). A similar behavior was observed in an EMSA titration for the corresponding DBD constructs (Fig. S3). ITC measurements allowed us to quantitatively analyze DNA binding thermodynamics and to precisely compare WT and variant constructs (Fig. 5C–E). Representative titration curves for DD-DBD/R263C and DD-DBD/N266S are shown in Fig. 5C,D. An average *K_D_* value of 7.9 μM was obtained for DD-DBD/R263C, which is significantly higher than the average *K_D_* value of 0.10 μM for the DD-DBD WT construct (Fig. 1H, Table 2). DBD/R263C exhibited slightly stronger binding than the dimeric construct, with an average *K_D_* value of 1.6 μM (Table 2). Similar to the WT proteins, the binding event for the R263C variants was an enthalpy-driven process (Fig. 5E). For the N266S variant, ITC experiments revealed average *K_D_* values of 0.57 μM (DBD/N266S) and 0.74 μM (DD-DBD/N266S), which again pointed towards an impaired protein-DNA interaction compared to WT proteins (Fig. 5D,E, Table 2). In addition, the stoichiometry dropped to an average number of approximately 1.6 (Table 2), which suggested a decrease in active protein concentration in the ITC cell. Indeed, a manual inspection of the sample after the ITC experiment confirmed that white precipitation had formed. As the ITC experiment was performed at 30 °C, this aggregation could be explained by the low thermal stability observed in DSF (Table 1).

**Fig. 5.**
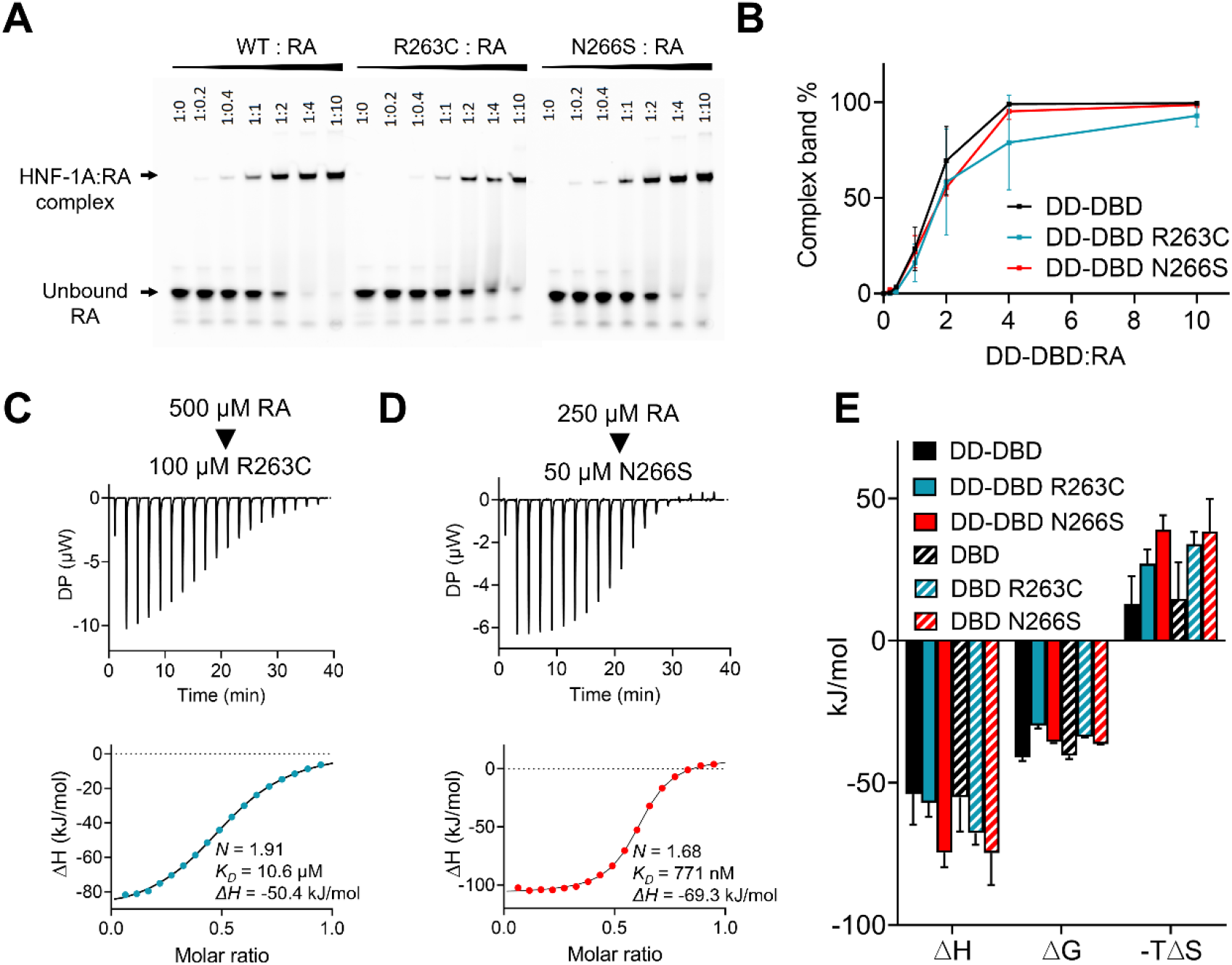
DNA binding parameters for HNF-1A R263C and N266S. A) EMSA gel for DD-DBD WT, R263C, and N266S variants. B) Quantification of EMSA gels for DD-DBD variants (N = 3). C, D) Representative ITC titration curve for DD-DBD/R263C (C) and DD-DBD/N266S (D). Titrant: 500μM (C) or 250 μM (D) double-stranded RA. Titrand: 100 μM DD-DBD/R263C (C) or 50 μM DD-DBD/N266S (D). E) Quantification of thermodynamic energies from ITC analysis (mean ± SD; N = 3).

**Table 2.**
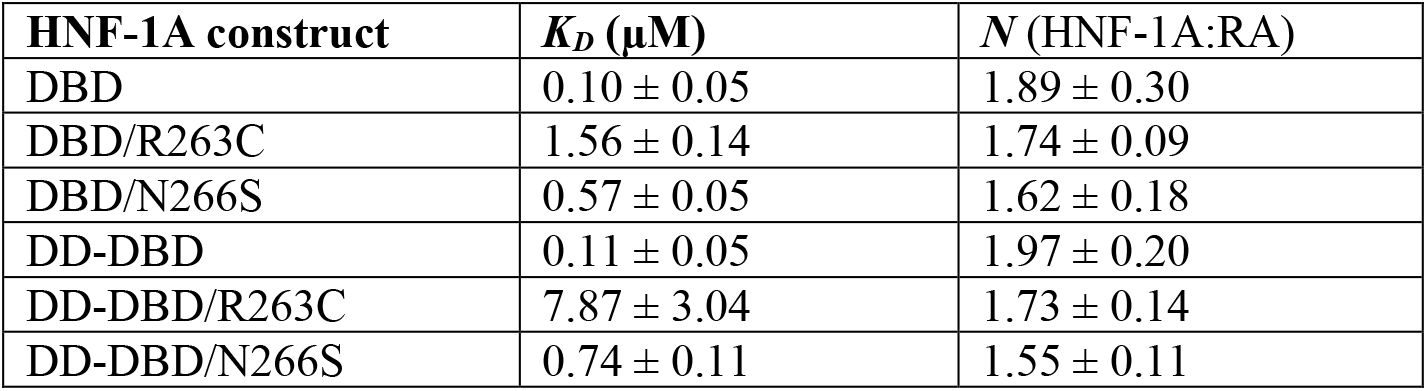
Protein:RA *K_D_* and *N* values determined by ITC (mean ± SD; N = 3). Parameters were extracted from binding experiments in which double-stranded RA oligonucleotide was titrated into DBD, DD-DBD, or corresponding HNF-1A variant constructs.

| HNF-1A construct | $K_D$ (μM) | $N$ (HNF-1A:RA) |
| --- | --- | --- |
| DBD | $0.10 \pm 0.05$ | $1.89 \pm 0.30$ |
| DBD/R263C | $1.56 \pm 0.14$ | $1.74 \pm 0.09$ |
| DBD/N266S | $0.57 \pm 0.05$ | $1.62 \pm 0.18$ |
| DD-DBD | $0.11 \pm 0.05$ | $1.97 \pm 0.20$ |
| DD-DBD/R263C | $7.87 \pm 3.04$ | $1.73 \pm 0.14$ |
| DD-DBD/N266S | $0.74 \pm 0.11$ | $1.55 \pm 0.11$ |

## Discussion

Considering the diverse expression patterns and major role of HNF-1A for a functional endocrine pancreas, remarkably little is known about the biochemical structure of HNF-1A and protein function on a molecular level. Notably, pathogenic HNF-1A variants cause the most common form of inherited diabetes, MODY3. Even though the genetic cause of MODY3 is known, there is limited molecular understanding of how a dysfunctional HNF-1A protein leads to decreased insulin secretion. We therefore aimed to extend the knowledge on the structural and biophysical characteristics of HNF-1A to study MODY3 variants in greater detail. In this work, we henceforth investigated two truncated versions of the HNF-1A protein and three MODY3-associated HNF-1A variants.

Our finding that DD-DBD can dimerize independently of the dimerization co-factors PCBD1 and PCBD2 was expected, as DD had been shown to form homodimers in isolation (8,9). Complex formation with PCBD1/2 may stabilize this dimeric form, as demonstrated by Bayle *et al*. (11), and thereby modulate HNF-1A function. Even though the DNA binding affinity was not significantly increased for the dimeric construct, the dimerization of HNF-1A may affect the protein’s lifetime by protecting it from denaturation and proteolysis (30), or could participate in transcriptional activity as dimerization may increase the local concentration of the regulatory TAD (31).

Moreover, we found that our HNF-1A constructs exhibit a high degree of flexibility. The 30-residue POU_S_-POU_H_ linker tethers the two homeodomains together and allows for distance variation between them. This intrinsic conformational flexibility is a general feature of the POU transcription factor family, enabling the proteins to adopt diverse quaternary structures during DNA recognition (32–34). Two well-characterized exemplary POU transcription factors are Oct-1 and Pit-1, which are both able to bind to DNA motifs in different arrangements, depending on the nature of the DNA response element (35). Oct-1 binds to both the *Palindromic Oct-factor response element* (PORE) as well as to *More of PORE* (MORE), albeit in different conformations (36) (Fig. 6). In the PORE binding mode, the POU_S_ and POU_H_ domains of one Oct-1 protomer are arranged on opposite sides of the DNA molecule, with the linker being extended across the double helix axis. In contrast, the POU_S_ and POU_H_ domains are in a compact formation on the same side of the DNA molecule when Oct-1 is bound to a MORE DNA motif (36). The DNA recognition mode of Oct-1 influences its ability to interact with the transcriptional enhancer OBF-1, which is only able to bind and activate Oct-1 transcriptional activity in the PORE-bound state (35,37,38). Thus, the conformational flexibility of the Oct-1 transcription factor is crucial for variable arrangements and gene activation modes. A similar conformational variability was shown for the POU transcription factor Pit-1 (39), where different DNA protein arrangements alter the accessibility for transcription modulators, such as the co-repressor N-CoR (35). These examples of protein-DNA recognition modes illustrate the importance of the flexible linker and allow speculations about HNF-1A not only recognizing the established promoters in a MORE-like fashion, but perhaps also binding to undescribed response elements with distinct HNF-1A promoter recognition modes.

**Fig. 6.**
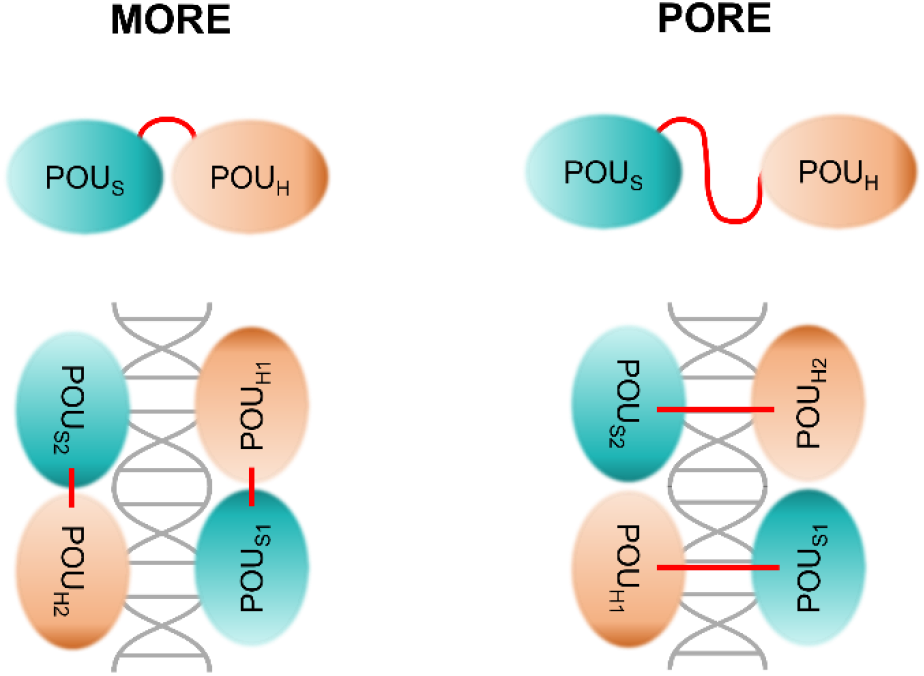
MORE and PORE DNA binding modes of POU transcription factors (32–36). Transcription factors in MORE arrangement are bound with both POU_S_ and POU_H_ domains on the same side of the DNA molecule. The homeodomains are in close proximity and the linker is compact. In PORE binding mode, the POU_S_ and POU_H_ domains are located on opposite sides of the DNA, with the linker being extended across the DNA symmetry axis.

The P112L variant has previously been functionally studied in cells (20,22). While the nuclear localization appears normal, HNF-1A P112L exhibits strongly impaired DNA binding ability *in vitro* and transactivation potential on the RA promoter in HeLa cells (20,22). Surprisingly, the P112L constructs were insoluble in our expression conditions. This insolubility could arise from structural instability, which can be explained by the substitution of Pro112 at the critical position in the beginning of helix α2 (Fig. 3B,C). Proline residues play important roles in modulating hydrogen bond formation in α-helix backbone structures (40–42), and the P112L substitution may lead to an N-terminally elongated α2 helix with impact on overall protein folding. Moreover, the rigidity and bulkiness of Pro112 may also contribute to protein folding by providing an optimal local backbone conformation.

The conducted protein turnover experiments in HeLa cells support our hypothesis of enhanced protein degradation dynamics. One way to further delineate the mechanism is the application of specific drugs that inhibit lysosomal and proteasomal degradation pathways. Our findings agree with a recent large-scale multidimensional analysis of MODY3 variants, where HNF-1A P112L expression in HeLa cells was lower compared to WT HNF-1A (43). Our data supports an alternative cause of disease for HNF-1A P112L, with P112L being a structurally destabilizing mutation, rather than a pure DNA binding mutation as suggested in earlier reports (20,22).

HNF-1A R263C proved to be a loss-of-function mutation regarding DNA binding. Our results are in accordance with earlier studies showing that *in vitro* translated HNF-1A R263C is unable to bind DNA and that this HNF-1A variant has a significantly decreased transactivation potential in HeLa cells (22,23). This effect can be explained by the positioning of Arg263 in HNF-1A (Fig. 3D). Arg263 undergoes direct contacts with o.A16 of the promoter motif and appears to form ion-dipole interactions with Leu258 and Thr260. Interestingly, a genetic variance in the latter (T260M) was identified in a MODY3 patient. It compromises DNA binding and causes impaired β-cell function (44). Based on these data, the Arg263-Thr260-Leu258 microenvironment and the direct interaction between Arg263 and o.A16 appear to be crucial for adequate HNF-1A DNA interaction (Fig. 3D).

Finally, we utilized this system to investigate an HNF-1A VUS (N266S), for which previously unknown mechanisms were uncovered. The mutant exhibits altered thermal stability, minor changes in folding, and a decrease in protein:DNA affinity. A combination of all effects may lead to impaired HNF-1A function in the pancreatic β-cell. Bioinformatic analyses of the protein region harboring the Asn266 residue may assist in explaining possible molecular mechanisms (Fig. S5). By performing a local alignment search for HNF-1A, we identified a homologous protein, Homeobox-containing protein 1 (HMBOX1) (45), which shares 28% sequence identity and exhibits a conserved homeobox domain (Fig. S5A). A structure alignment between the HNF-1A DBD crystal structure (16) and the crystal structure of the HMBOX1 homeodomain (45) showed high structural similarity between the compared domains (Fig. S5B). Especially the amino acid sequence and structural arrangement of helix α9 is highly conserved between the two proteins (Fig. S5A,B), indicating that residues in this region are critical for HNF-1A and HMBOX1 function. Asn266 in HNF-1A may form polar interactions important for the stabilization of the α-helical backbone structure of α9 or might indirectly bind to the DNA. Hence, the N266S substitution may compromise helix stability or DNA binding. Alternatively, the substitution of Asn266 with Ser may potentially introduce a phosphorylation site, which could de-stabilize protein structure and hinder DNA binding. Interestingly, HMBOX1 is strongly expressed in pancreatic tissue, where it may act as a transcriptional repressor (46). The conservation of the homeodomain and the α9 helix in HNF-1A and HMBOX1 allow speculations that both proteins are able to bind to the same target promoter sequence; HNF-1A as a transcriptional activator, and HMBOX1 as a transcriptional repressor. Due to compromised α9 integrity and DNA binding, the N266S variant may be outcompeted by HMBOX1, leading to an altered regulation of gene transcription in β-cells. Further studies in human cell lines will likely complete our understanding of disease causality, possibly leading to a re-classification of HNF-1A N266S from VUS to (likely) pathogenic.

In summary, we established a biophysical characterization platform for HNF-1A, which proved to be a valuable tool to investigate MODY3 mutations in the DD and DBD of HNF-1A. We verified pathogenic effects, as well as provided further understanding for structural and functional dysregulation of the variants. In particular, primary effects caused by structurally de-stabilizing mutations may be uncovered in a direct manner, providing explanations for secondary effects in gene transactivation activity.

## Conclusions

Structural and biochemical studies is an underrepresented approach in the variant interpretation process of transcription factor-associated diseases, such as MODY. Our presented methodology provides new insights into WT HNF-1A protein function, and demonstrates that structure-based investigations may lead to increased certainty in variant-phenotype correlation, and can play a valuable part in MODY precision diagnostics and treatment.

## Experimental procedures

### Plasmid generation

The human *HNF1A* gene in a pcDNA™3.1/HisC plasmid (22) was used in order to subclone two HNF-1A constructs (DD-DBD: residues 1 - 279, DBD: residues 83 - 279) into the Gateway donor vector pDONR221 (Invitrogen™). An N-terminal Tobacco-Etch Virus (TEV) protease site (ENLYFQG) was included. In addition, attB1 and attB2 sites were introduced N- and C-terminally of the respective gene fragments, allowing for Gateway cloning strategy. Final expression clones in the destination vector pTH27 (47) harbored an N-terminal His_6_-tag (His_6_-(DD-)DBD). MODY3-associated variants (P112L, R263C, N266S) were introduced into (DD-)DBD pTH27 expression plasmids by Q5® site-directed mutagenesis (New England Biolabs, Ipswich, MA, USA). These constructs were used for recombinant protein expression and purification, in order to biophysically and functionally characterize HNF-1A variants.

Using the same strategy, the full-length *HNF1A* gene sequence (residues 1 - 631) was subcloned into the pcDNA™3.1/nV5-DEST mammalian expression vector (Invitrogen™), harboring an N-terminal V5-tag. The P112L variant was generated by Q5® site-directed mutagenesis (New England Biolabs, Ipswich, MA, USA). These constructs were used to assess protein stability in a CHX assay.

Primers used in Gateway cloning, Q5^®^ site-directed mutagenesis, and plasmid sequencing are listed in table S4.

### Recombinant protein expression and purification

His_6_-(DD-)DBD constructs were expressed in the *E. coli* Rosetta (DE3). Bacteria were grown in *Luria-Bertani* (LB) medium at 37 °C until an OD_600_ value of 0.6 - 0.8 was reached. Protein expression, induced by the addition of 1 mM of isopropyl β-d-1-thiogalactopyranoside (IPTG), was performed at 20 °C for 20 h.

Bacterial cell pellets were resuspended in 50 mM HEPES (pH 7.5), 500 mM NaCl, 1 mM dithiothreitol (DTT), 20 mM imidazole, 1x cOmplete™ EDTA-free protease inhibitor cocktail (Roche, Basel, Switzerand), 1 mM phenylmethylsulphonyl fluoride (PMSF), and lyzed by ultrasonication (7 min, 1 s on/off cycles, 25 W). Cell lysates were cleared by centrifugation (16 000 × *g*, 30 min, 4 °C) and 6xHis-(DD-)DBD was purified using Ni-NTA affinity chromatography. Elution was done using an imidazole gradient (40 – 300 mM) and protein containing fractions were dialyzed overnight at 4 °C (20 mM HEPES pH 7.5, 500 mM NaCl, 1 mM DTT). The N-terminal His_6_-tags were proteolytically removed by the addition of TEV protease during the dialysis step. A second Ni-NTA affinity chromatography step was used to separate cleaved from uncleaved proteins, where a gradient in a low-imidazole range was applied (0 mM – 80 mM). Relevant fractions were concentrated and subjected to SEC, using a Superdex 75 pg 16/60 column (GE Healthcare, Chicago, IL, USA). The running buffer contained 20 mM HEPES pH 7.5, 500 mM NaCl, and 1 mM TCEP. Fractions containing pure (DD-)DBD were concentrated, snap-frozen in liquid N2, and stored at −80 °C.

The expression and solubility of mutant constructs was assessed in *E. coli* Rosetta (DE3) and *E. coli* Lemo21 (DE3) strains, varying expression temperatures and lysis buffer composition. HNF-1A variants R263C and N266S were recombinantly expressed and purified in the same way as WT proteins. The HNF-1A P112L variant was not expressed and purified due to insolubility in the tested conditions.

Protein identity was confirmed by peptide mass fingerprinting using a Bruker Ultra fleXtreme MALDI-TOF mass analyzer.

### Dynamic light scattering

Monodispersity and R_h_ of (DD-)DBD variants was assessed using DLS. Measurements of 1.5 mg/ml purified variants (20 mM HEPES pH 7.5, 500 mM NaCl, 1 mM TCEP) were performed using a Malvern Zetasizer Nano-ZS instrument at 25 °C. Each variant was measured in triplicate.

### Size exclusion chromatography – multi-angle light scattering

Molecular weights of (DD-)DBD and respective variants were determined using SEC-MALS. Chromatography was performed at room temperature, on a Superdex 200 Increase 10/300GL (GE Healthcare, Chicago, IL, USA) column in SEC buffer (20 mM HEPES pH 7.5, 500 mM NaCl, 1 mM TCEP). 100 μg sample was subjected to gel electrophoresis at a flow rate of 0.5 ml/min and MALS was recorded using a miniDAWN TREOS instrument. Concentration determination was based on refractive index, recorded with an on-line ERC RefractoMax 520 refractometer. The monomeric third fibronectin III domain of rat β4-integrin (MW = 12.76 kDa) was used as internal molecular weight standard (48). Data were analyzed using the ASTRA software (Wyatt, Santa Barbara, CA, USA).

### Differential scanning fluorimetry

DSF measurements were performed using a Lightcycler 480 II (Roche, Basel, Switzerland) thermocycler. 0.25 mg/ml DD-DBD or 0.5 mg/ml DBD variants in 2 mM HEPES pH 7.5, 50 mM NaCl, 0.1 mM TCEP, 15x SyproOrange (Invitrogen™) were subjected to thermal denaturation (20 to 95 °C, 0.2 °C data pitch). SyproOrange fluorescence at 465-610 nm was followed. All measurements were performed as technical triplicate. The fluorescence signal was normalized and plotted against temperature. Graphpad Prism 8.3.0 (GraphPad Software, San Diego, CA, USA) was used to fit the data to a sigmoidal function and to extract *T_M_* values from the inflection point of the curve.

### Small-angle X-ray scattering

Initial SAXS experiments on WT proteins were performed on the P12 synchrotron beamline (49), EMBL/DESY (Hamburg, Germany), as mail-in service. Batch measurements were done in 20 mM HEPES (pH 8.0), 500 mM NaCl, 1 mM TCEP at 10 °C. A dilution series (0.5 mg/ml – 2.0 mg/ml) was prepared in order to investigate potential concentration-dependent effects. Bovine serum albumin was used as standard protein for molecular weight determination.

SAXS measurements of all variants were performed as mail-in service on the SWING beamline (50) at the SOLEIL synchrotron (Saint-Aubin, France). All measurements were performed in SEC-SAXS mode using an Agilent Biosec3-300 column at 14 °C (20 mM HEPES pH 7.5, 500 mM NaCl, 2 mM TCEP). 50 μl of protein sample at 4 – 7 mg/ml was injected, and X-ray scattering data were collected as the sample eluted from the column.

SAXS data were reduced and processed using the ATSAS 3.0.3 package (51). CRYSOL (52) was employed to generate a theoretical scattering curve of the DNA-bound DBD from the published DBD:DNA crystal structure (PDB: 1IC8) (16), which was subsequently compared to the experimental scattering curve of the DNA-free DBD in solution. *Ab initio* models for (DD-)DBD were generated by using GASBOR (53). Rigid body models of multi-domain proteins were generated by using CORAL (54). Here, the isolated POU_S_ and POU_H_ domains (PDB: 1IC8) and the dimeric DD (PDB: 1F93, (10)) were used as rigid bodies, and linker regions were modelled as flexible loops. Further modelling details are listed in table S2 and S3. Visualization of SAXS models was done in PyMOL (55).

### Electrophoretic mobility shift assay

EMSA was used to qualitatively assess the DNA binding ability of HNF-1A variants. Cy5-labelled oligonucleotide probes, corresponding to the RA promoter as found in PDB 1IC8 (16), were commercially synthesized (Merck KGaA, Darmstadt, Germany) and dissolved in 10 mM Tris pH 8.0, 50 mM NaCl, 1 mM EDTA to 100 μM final concentration. 5’ oligo ([Cy5]-CTTGGTTAATAATTCACCAGA) and 3’ oligo ([Cy5]-TCTGGTGAATTATTAACCAAG) were mixed in an equimolar ratio. The probes were annealed into double-stranded DNA by incubation at 95 °C for 5 min, before being allowed to cool down to room temperature over 45 min.

EMSA was performed using the LI-COR EMSA buffer kit (LI-COR Biosciences, Lincoln, NE, USA). The EMSA reaction mix contained 1x Binding buffer (10 mM Tris pH 7.5, 50 mM KCl, 1 mM DTT), 0.5 μM [Cy5]-RA, 2 mM DTT, and varying amounts of protein sample. The DNA probe was mixed in different ratios with purified (DD-)DBD variants, sampling DNA:protein molar ratios from 1:0 to 1:10. EMSA was performed according to protocol instructions and Cy5 signal was detected from the EMSA gel using a ChemiDoc MP imaging system.

### Isothermal titration calorimetry

ITC measurements were performed using a MicroCal iTC200 instrument (Malvern Panalytical, Malvern, UK). Unlabelled oligonucleotides (TAG Copenhagen, Copenhagen, Denmark) were annealed into double-stranded RA as described above. (DD-)DBD variants and the DNA were dialyzed against 20 mM HEPES pH 7.5, 200 mM NaCl, 1 mM TCEP. Typically, 50 μM RA was titrated into 10 μM (DD-)DBD protein. Higher concentrations were required for the R263C and N266S variants, where 250 μM RA was titrated into 50 μM protein. The injection program contained 19 injections with 2 μl injection volume, with a two-minute spacing interval. All measurements were performed at 30 °C in triplicate from the same protein batch. Titrations were analyzed using the MicroCal PEAQ-ITC analysis software (Malvern Panalytical, Malvern, UK). *ΔH, ΔG, TΔS, K_D_*, and *N* were extracted from the fits.

### Synchrotron radiation circular dichroism

SRCD measurements were performed on the AU-CD beamline, ASTRID2 synchrotron, ISA, University of Aarhus, Denmark. Purified proteins and double-stranded RA were dialyzed into CD buffer (20 mM Na-phosphate buffer pH 8.0, 150 mM NaF, 1 mM TCEP). Initial measurements were conducted at 25 °C for WT constructs and HNF-1A:RA complex formation studies. SRCD measurements for the comparison of WT HNF-1A and MODY3 variants were performed at 10 °C. 0.1-mm circular cells (Hellma Analytics, Müllheim, Germany) were used. Initial SRCD spectra on WT constructs and HNF-1A:RA complexes were recorded in the wavelength range of 350 – 170 nm, while spectra for WT-variant comparisons were recorded in a wavelength range of 280 – 170 nm. Wavelength scans were performed in 1 nm increments. Three scans per sample were performed and averaged. Generally, protein constructs were measured at a concentration of 1 mg/ml. For measurements of protein:DNA complexes, double-stranded RA oligonucleotide and (DD-)DBD constructs were mixed in a 1:2 molar ratio. Data processing was done using CDtoolX (56) or using Microsoft Excel.

### Cycloheximide chase assay

HeLa cells were grown in DMEM growth medium (Sigma-Aldrich, St. Louis, MO, USA), supplemented with 15% fetal bovine serum, 1% penicillin-streptomycin, and 4 mM L-glutamine, in a humidity incubator keeping constant conditions of 37 °C and 5% atmospheric CO_2_. HeLa cells were seeded in 6-well plates at a density of 200 000 cells per well. 4 × 4 wells were prepared for each variant (WT / P112L HNF-1A). Transfection with full-length nV5-WT or nV5-P112L (0.8 μg DNA/well) was performed 24 h post-seeding using X-tremeGENE™ 9 DNA Transfection Reagent (Roche, Basel, Swityerland). 24 h post-transfection, the growth medium was exchanged with growth medium supplemented with 50 μg/ml CHX (Merck KGaA, Darmstadt, Germany). Sample acquisition was done at different time points of CHX treatment (0 h, 2 h, 4 h, 6 h). HeLa cells were washed twice with ice-cold phosphate-buffered saline (PBS) and harvested in 1 ml ice-cold PBS using a cell scraper. Samples were centrifuged (16 000 x *g*, 4 °C, 2 × 15 s) and the dry cell pellets were stored at −80 °C. Cells were resuspended in 40 μl lysis buffer (50 mM Tris pH 8.0, 150 mM NaCl, 5 mM EDTA, 1% NP-40, 1x cOmplete™ EDTA-free protease inhibitor cocktail) and incubated on ice for 30 min. Cell lysates were cleared by centrifugation (16 000 x *g*, 10 min, 4 °C) and concentrations of the soluble fraction were determined by using the Pierce™ BCA protein assay (Thermo Fisher Scientific, Waltham, MA, USA). 20 μg protein samples were analyzed by SDS-PAGE and Western blotting. Following protein transfer, the membrane was blocked in 5% milk-PBS-Tween (milk-PBS-T). Primary antibodies anti-V5 (Invitrogen, R960-25) and anti-pan-actin (Abcam, ab200658) were applied at 1:2000 in 1% milk-PBS-T and incubated overnight at 4 °C. Membranes were washed three times with PBS-T. Secondary antibody was applied at a 1:5000 dilution in 3% milk-PBS-T and incubated for 2 h at 4 °C. Membranes were washed three times in PBS-T before horseradish peroxidase (HRP) signals were developed with SuperSignal® West Pico Chemiluminescent Substrate (Thermo Fisher Scientific, Waltham, MA, USA). Detection of HRP signal was conducted using a ChemiDoc™ XRS+ imaging system. Band intensities were quantified using the ImageLab™ v3.0 software (Bio-Rad Laboratories, Hercules, CA, USA). V5-bands, representing protein levels of V5-WT and V5-P112L variants, were normalized to total protein amount based on stain-free gel imaging prior to Western blot transfer (Fig. S2). Signals for V5-WT and V5-P112L at different time points of CHX treatment were further normalized to the respective V5-signal at 0 h treatment. Anti-actin bands were used to visualize equal protein loading. Statistical analysis was done in GraphPad Prism version 8.3.0., using an unpaired t-test per time-point without assuming a consistent SD. Statistical significance was determined using the Holm-Sidak method, with a significance threshold of 0.05.

### Bioinformatic analysis

Protein disorder prediction was done using the IUPred2 prediction server (26). Prediction score curves produced by IUPred2A and DynaMine (27) algorithms were plotted in Graphpad Prism 8.3.0.

Sequence conservation of the HNF-1A DBD was investigated by the generation of a global alignment. HNF-1A protein sequences from several model organisms were extracted from the UniProt (57) database (*Homo sapiens*: P20823, *Mus musculus*: P22361, *Danio rerio*: Q4L1M9, *Xenopus laevis*: Q05041, *Gallus gallus*: Q90867). Sequences were aligned using Clustal Omega (58) and a graphical presentation was generated using ESPript 3.0 (59).

Thermal stability of HNF-1A variants was predicted by using the CUPSAT algorithm (29), considering torsion angle distributions and amino acid atom potentials. As convention, a negative ΔΔG corresponds to a de-stabilizing point mutation.

A local alignment search for HNF-1A was conducted using the Basic Local Alignment Search Tool (BLAST) (60). A pair-wise alignment of human HNF-1A (UniProt ID: P20823) and human HMBOX1 (UniProt ID: Q6NT76) was generated using Clustal Omega (58) and a graphical presentation was generated using ESPript 3.0 (59). A structure alignment of the DBD of HNF-1A (PDB: 1IC8, chain A, (16)) and the homeobox domain of HMBOX1 (PDB: 4J19, chain A, (45)) was generated with the RCSB PDB pair-wise structure alignment tool using the jFATCAT rigid algorithm (61). Protein structures were analyzed and visualized using PyMOL (55).

## Acknowledgements

We acknowledge Rezan Erman and Marcus Langeland Larsen Nygård for technical assistance. We are thankful for guidance with SEC-MALS measurements by Dr. Anne Baumann. ITC, DSF, DLS, and SEC-MALS experiments were performed at the Biophysics, Structural Biology and Screening (BiSS) facility at the University of Bergen. We thank Ulrich Bergmann (Biocenter Oulu Proteomics Core Facility) for conducting mass spectrometry analysis. We are grateful for beamtime and supportive synchrotron beamline staff at PETRAIII EMBL P12, SOLEIL Synchrotron SWING, and ASTRID2 ISA AU-CD beamlines.

## Funding and additional information

This work was funded with a PhD fellowship by the Medical Faculty, University of Bergen, UiB, Norway (to L.K.), and supported by a UiB Meltzer foundation project grant (to L.K.). L.K. received a research visit grant from the Norwegian Graduate School in Biocatalysis. T.A. was supported by the Norwegian Cancer Society (Project 171752—PR-2009-0222). P.R.N. received funding from the European Research Council (AdG SELECTionPREDISPOSED #293574), the Research Council of Norway (FRIPRO grant #240413), the Western Norway Regional Health Authority (Strategic Fund “Personalized Medicine for Children and Adults”), the Novo Nordisk Foundation (grant #54741), and the Norwegian Diabetes Association. This work has been supported by the project CALIPSOplus under the Grant Agreement 730872 from the EU Framework Programme for Research and Innovation HORIZON 2020 (to P.K.).

