## Supporting information for "Structural and biophysical characterization of HNF-1A as a tool to study MODY3 diabetes variants"

### List of included material

Table S1**.** Molecular weight of DBD and DD-DBD, determined by SEC-MALS.

Table S2. Small-angle X-ray scattering parameters for DBD and DD-DBD WT constructs.

Table S3. Small-angle X-ray scattering parameters for HNF-1A variant constructs.

Table S4. Primers used in Gateway cloning, Q5 mutagenesis, and gene insert sequencing.

Fig. S1. WT/P112L expression tests in *E. coli* Rosetta(DE3) and Lemo21(DE3).

Fig. S2. Protein turnover dynamics experiment using CHX assay.

Fig. S3. Representative EMSA gels for DBD WT, R263C, and N266S variants.

Fig. S4. Thermal stability of WT, R263C, and N266S variants.

Fig. S5. Sequence and structure alignment of HNF-1A and HMBOX1.

**Table S1. Molecular weight of DBD and DD-DBD, determined by SEC-MALS.**

| Variant | DD-DBD ($M_{monomeric}$ = 31.2 kDa) | DBD ($M_{monomeric}$ = 22.8 kDa) |
|---|---|---|
| WT | 61.3 kDa (dimer) | 24.0 kDa (monomer) |
| R263C | 60.2 kDa (dimer) | 23.9 kDa (monomer) |
| N266S | 61.9 kDa (dimer) | 24.1 kDa (monomer) |

**Table S2. Small-angle X-ray scattering parameters for DBD and DD-DBD WT constructs.**

| | DBD | DD-DBD |
|---|---|---|
| **Data collection** | | |
| Instrument | P12, PETRA III, DESY, Hamburg | |
| Detector | Pilatus 6M | |
| Wavelength (nm) | 0.124 | |
| Angular range ($nm^{-1}$) | 0.03 – 7.4 | |
| Temperature (°C) | 10 | |
| SAXS mode | Batch | |
| Exposure time (s) | 0.045 | |
| Concentration (mg/ml) | 0.5 – 2.0 | |
| **Software** | | |
| Primary data reduction and processing | PRIMUS | |
| Data validation and analysis | PRIMUS | |
| Generation and fitting of theoretical scattering profiles | CRYSOL | |
| *Ab initio* modelling | GASBOR | |
| Rigid body modelling | CORAL | |
| Graphical representation | PyMOL | |
| **Structural parameters** | | |
| $I_0$ (relative), from Guinier | 143.9 | 598.0 |
| $R_g$ (nm), from Guinier | 2.68 | 5.51 |
| $I_0$ (relative), from P(r) | 145.4 | 615.9 |
| $R_g$ (nm), from P(r) | 2.83 | 6.07 |
| $D_{max}$ (nm), from P(r) | 10.2 | 24.4 |
| Quality estimate | 0.71 | 0.70 |
| **Molecular weight determination** | | |
| M (kDa), theoretical from sequence | 22.78 | 31.21 |
| M (kDa), from $I_0$ | 14.14 | 58.74 |
| ***Ab initio* modelling** | | |
| Symmetry | P1 | P2 |
| $\chi^2$ | 1.07 | 1.55 |
| **Rigid body modelling** | | |
| Crystal structure | 1IC8 | 1IC8, 1F93 |
| Flexible residues in HNF-1A | 181 - 208 | 31 – 92, 181 - 208 |
| Symmetry | P1 | P1 |
| $\chi^2$ | 1.42 | 1.66 |

**Table S3. Small-angle X-ray scattering parameters for HNF-1A variant constructs.**

| | **DBD/WT** | **DBD/R263C** | **DBD/N266S** | **DD-DBD/WT** | **DD-DBD/R263C** | **DD-DBD/N266S** |
|---|---|---|---|---|---|---|
| **Data collection** | | | | | | |
| Instrument | SWING, SOLEIL, Paris, France | | | | | |
| Detector | EIGER-4M | | | | | |
| Wavelength (nm) | 1.03 | | | | | |
| Angular range ($nm^{-1}$) | 0.04 – 5.7 | | | | | |
| Temperature (°C) | 14 | | | | | |
| SAXS mode | SEC-SAXS (Biosec3-300 column) | | | | | |
| Injection volume (µl) | 50 | | | | | |
| Exposure time (s/frame) | 0.99 | | | | | |
| Concentration (mg/ml) | 5.2 | 5.8 | 4.3 | 5.0 | 6.4 | 6.7 |
| **Software** | | | | | | |
| Primary data reduction and processing | PRIMUS | | | | | |
| Data validation and analysis | PRIMUS | | | | | |
| **Structural parameters** | | | | | | |
| $I_0$ (relative), from GNOM | 7.10E-03 | 7.08E-03 | 7.16E-03 | 1.60E-02 | 1.70E-02 | 1.67E-02 |
| $I_0$ (relative), from Guinier | 7.14E-03 | 7.17E-03 | 7.27E-03 | 1.64E-02 | 1.69E-02 | 1.74E-02 |
| $I_0$ (relative), from P(r) | 7.14E-03 | 7.17E-03 | 7.27E-03 | 1.64E-02 | 1.69E-02 | 1.74E-02 |
| $R_g$ (nm), from GNOM | 2.65 | 2.64 | 2.75 | 5.18 | 5.28 | 5.24 |
| $R_g$ (nm), from Guinier | 2.79 | 2.86 | 2.83 | 5.65 | 5.69 | 5.77 |
| $R_g$ (nm), from P(r) | 2.79 | 2.86 | 2.84 | 5.67 | 5.70 | 5.80 |
| $D_{max}$ (nm), from P(r) | 10.0 | 10.5 | 11.5 | 2.4 | 2.2 | 2.4 |
| Quality estimate | 0.77 | 0.74 | 0.67 | 0.66 | 0.59 | 0.60 |
| **Molecular weight determination** | | | | | | |
| Overall – Bayesian (kDa) | 23.1 | 23.7 | 21.2 | 72.4 | 78.5 | 62.4 |

**Table S4. Primers used in Gateway cloning, Q5 mutagenesis, and gene insert sequencing.** Variants are described using HGVS nomenclature (https://varnomen.hgvs.org/) based on the following NCBI reference sequences (RefSeq): *HNF1A* NM_000545.8.

| Primer | Forward/ Reverse | Construct | Purpose |
|---|---|---|---|
| **Gateway cloning** | | | |
| TCTGAGAATCTTTATTTTCAGGGC ATGGTTTCTAAACTGAGCCAGCTG | Forward | DD-DBD, DD-DBD-TAD | Introduction of TEV-site and partial attB1 site |
| TCTGAGAATCTTTATTTTCAGGGC CCACCCATCCTCAAAGAGCTGG | Forward | DBD | Introduction of TEV-site and partial attB1 site |
| AGAAAGCTGGGTCTTACTGGG AGGAAGAGGCCATCTGG | Reverse | DD-DBD-TAD | Introduction of partial attB2 site |
| AGAAAGCTGGGTCTTAGTGCCG GAAGGCTTCTTCTTTG | Reverse | DBD, DD-DBD | Introduction of partial attB2 site |
| GGGGACAAGTTTGTACAAAAAA GCAGGCTCTGAGAATC | Forward | DBD, DD-DBD, DD-DBD-TAD | Completion of attB1 site |
| GGGGACCACTTTGTACAAGAAAG CTGGGT | Reverse | DBD, DD-DBD, DD-DBD-TAD | Completion of attB1 site |
| **Q5 mutagenesis** | | | |
| GCAGGAGGACctgTGGCGTGTGG | Forward | DBD, DD-DBD, DD-DBD-TAD | c.335C>T p.Pro112Leu mutagenesis |
| AGAAGGGTCTCCACCACGGC | Reverse | DBD, DD-DBD, DD-DBD-TAD | c.335C>T p.Pro112Leu mutagenesis |
| CGTGTCTACAgCTGGTTTGCC | Forward | DBD, DD-DBD | c.797A>G p.Asn266Ser mutagenesis |
| CACCTCCGTGACGAGGTT | Reverse | DBD, DD-DBD | c.797A>G p.Asn266Ser mutagenesis |
| GGAGGTGtGTGTCTACAACTGG | Forward | DBD, DD-DBD | c.787C>T p.Arg263Cys mutagenesis |
| GTGACGAGGTTGGAGCC | Reverse | DBD, DD-DBD | c.787C>T p.Arg263Cys mutagenesis |
| **Insert sequencing** | | | |
| TAATACGACTCACTATAGGG | Forward | DBD, DD-DBD, DD-DBD-TAD | Sequencing from T7 promoter (pTH27, pcDNA3.1-nV5-DEST) |
| TAGAAGGCACAGTCGAGG | Reverse | DD-DBD-TAD | Sequencing from BGH PolyA motif (pcDNA3.1-nV5-DEST) |
| GCTAGTTATTGCTCAGCGG | Reverse | DBD, DD-DBD | Sequencing from T7 terminator motif (pTH27) |
| GAAGAACCCTAGCAAGG | Forward | DD-DBD-TAD | Sequencing from *HNF1A* c.663 |
| GCCCGATGGTCATGAC | Reverse | DD-DBD-TAD | Sequencing from *HNF1A* c.1246 |

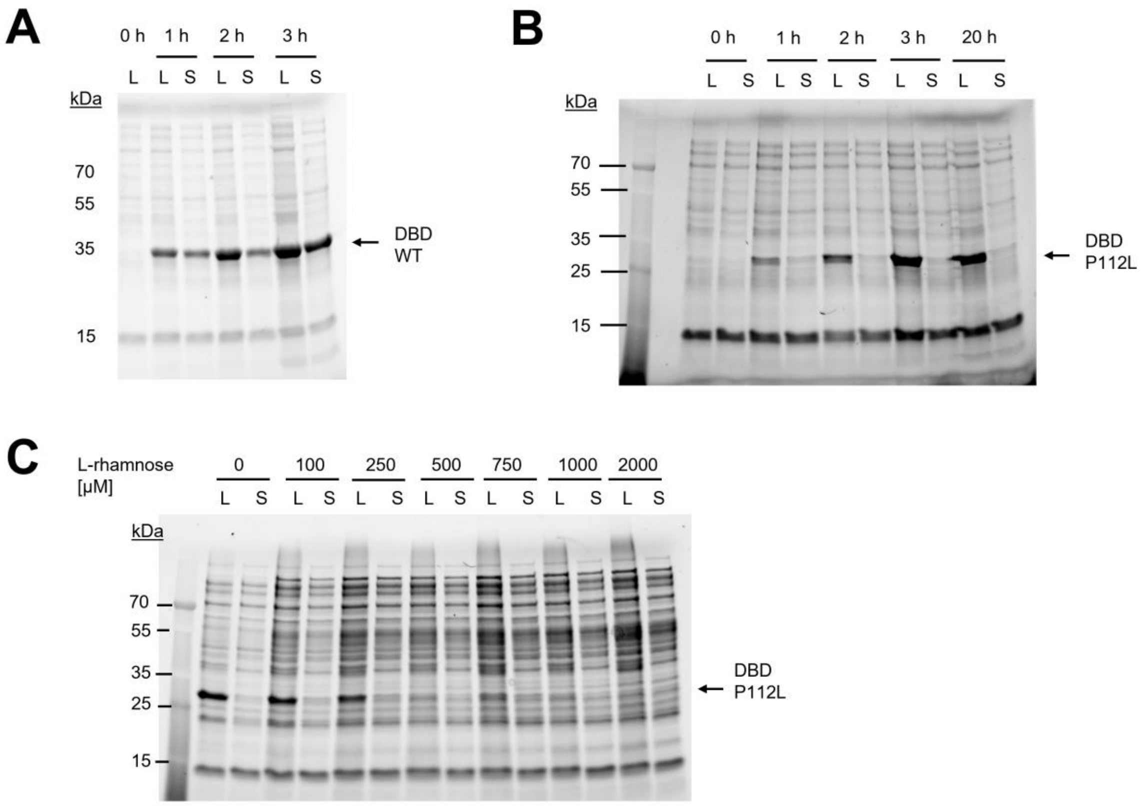


**Fig. S1. WT/P112L expression tests in *E. coli* Rosetta(DE3) and Lemo21(DE3).** A) Expression and solubility test for DBD WT in *E. coli* Rosetta(DE3), assessed by stain-free SDS-PAGE analysis. L – lysate, S – soluble fraction. 0 h – uninduced control. 1 – 3 h – expression time at 37 °C. B) Expression and solubility test for DBD/P112L in *E. coli* Rosetta(DE3), assessed by stain-free SDS-PAGE analysis. L – lysate, S – soluble fraction. 0 h – uninduced control. 1 – 3 h and 20 h – expression time at 37 °C and 20 °C, respectively. C) Expression and solubility test for DBD/P112L in *E. coli* Lemo21(DE3), assessed by stain-free SDS-PAGE analysis. L – lysate, S – soluble fraction. 5 h expression at 30 °C. Tunable expression experiment, performed with L-rhamnose titration.

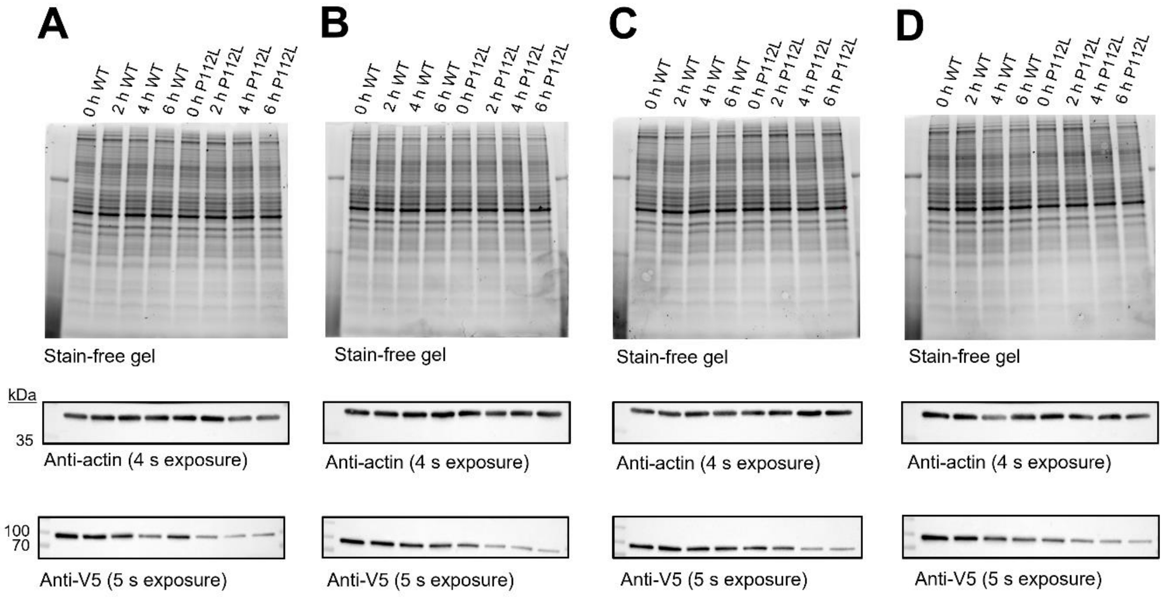


**Fig. S2. Protein turnover dynamics experiment using CHX assay.** GelDoc stain-free images and ChemiDoc western blot membranes for the CHX assay, assessing protein turnover dynamics of V5-HNF1A-WT and V5-HNF1A-P112L full-length proteins. A – D) Technical replicates 1 – 4.

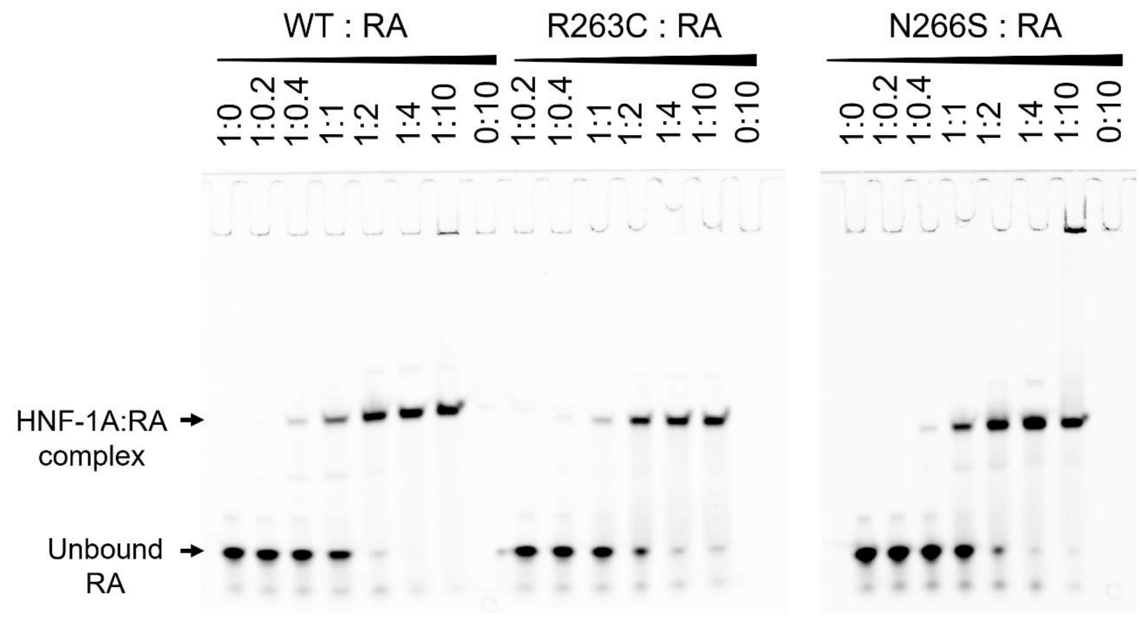


**Fig. S3. Representative EMSA gels for DBD WT, R263C, and N266S variants.**

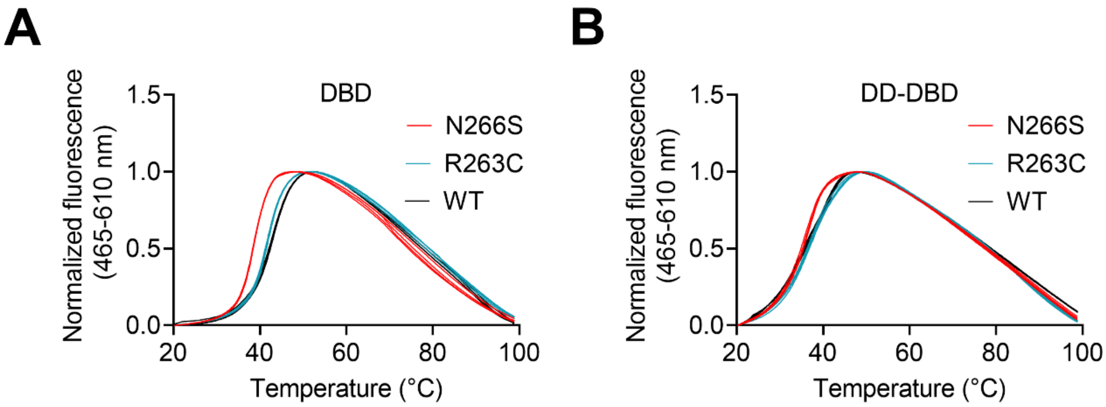


**Fig. S4. Thermal stability of WT, R263C, and N266S variants.** DSF melting curves for DBD (A) and DD-DBD (B) constructs (N = 4). One representative melting curve *per* construct.

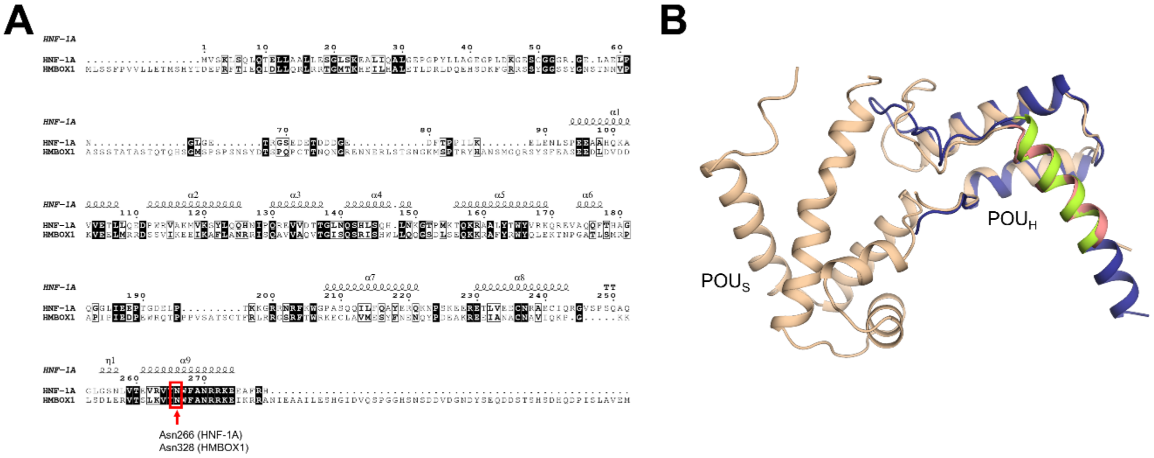


**Fig. S5. Sequence and structure alignment of HNF-1A and HMBOX1.** A) Local sequence alignment of HNF-1A and HMBOX1. Conserved residues are marked with dark background, and residues of interest (Asn266 in HNF-1A, Asn328 in HMBOX1) are indicated. B) Structure alignment of the HNF-1A DBD crystal structure (PDB: 1IC8, wheat) and the crystal structure of the homeobox domain of HMBOX1 (PDB: 4J19, blue). Conserved helix α9 is colored in salmon and green for HNF-1A and HMBOX1, respectively.
